# Genomic and chemical diversity of *Bacillus subtilis* secondary metabolites against plant pathogenic fungi

**DOI:** 10.1101/2020.08.05.238063

**Authors:** Heiko T. Kiesewalter, Carlos N. Lozano-Andrade, Mario Wibowo, Mikael L. Strube, Gergely Maróti, Dan Snyder, Tue Sparholt Jørgensen, Thomas O. Larsen, Vaughn S. Cooper, Tilmann Weber, Ákos T. Kovács

## Abstract

*Bacillus subtilis* produces a wide range of secondary metabolites providing diverse plant-growth-promoting and biocontrol abilities. These secondary metabolites include non-ribosomal peptides (NRPs) with strong antimicrobial properties, causing either cell lysis, pore formation in fungal membranes, inhibition of certain enzymes, or bacterial protein synthesis. However, the natural products of *B. subtilis* are mostly studied either in laboratory strains or in individual isolates and therefore, a comparative overview of *B. subtilis* secondary metabolites is missing.

In this study, we have isolated 23 *B. subtilis* strains from eleven sampling sites, compared the fungal inhibition profiles of wild types and their NRPs mutants, followed the production of targeted lipopeptides, and determined the complete genomes of 13 soil isolates. We discovered that non-ribosomal peptide production varied among *B. subtilis* strains co-isolated from the same soil samples. *In vitro* antagonism assays revealed that biocontrol properties depend on the targeted plant pathogenic fungus and the tested *B. subtilis* isolate. While plipastatin alone is sufficient to inhibit *Fusarium* sp., a combination of plipastatin and surfactin is required to hinder the growth of *Botrytis cinerea*. Detailed genomic analysis revealed that altered NRP production profiles in certain isolates is due to missing core genes, nonsense mutation, or potentially altered gene regulation.

Our study combines microbiological antagonism assays with chemical NRPs detection and biosynthetic gene cluster predictions in diverse *B. subtilis* soil isolates to provide a broader overview of the secondary metabolite chemodiversity of *B. subtilis*.

**IMPORTANCE:** Secondary or specialized metabolites with antimicrobial activities define the biocontrol properties of microorganisms. Members of the *Bacillus* genus produce a plethora of secondary metabolites, of which non-ribosomally produced lipopeptides in particular display strong antifungal activity. To facilitate prediction of the biocontrol potential of new *Bacillus subtilis* isolates, we have explored the *in vitro* antifungal inhibitory profiles of recent *B. subtilis* isolates, combined with analytical natural product chemistry, mutational analysis, and detailed genome analysis of biosynthetic gene clusters. Such a comparative analysis helped to explain why selected *B. subtilis* isolates lack production of certain secondary metabolites.

## INTRODUCTION

The rhizosphere is well known as a microbial hotspot since it can be seen as a nutrient-rich oasis surrounded by otherwise nutrient-limited regions of soil. This ecosystem comprises a plethora of intra- and interspecies interactions between bacteria, fungi, plants and higher organisms mediated by a diversity of natural products. Soil bacteria, in general, are capable of producing a huge amount of different secondary or specialized metabolites, which although not essential for growth, might have miscellaneous functions. However, our understanding of the true ecological role of these specialized metabolites has just begun to unfold. On the one hand, secondary metabolites are assumed to be biological weapons that provide the producer strains a competitive advantage in asserting themselves in an ecological niche (1). On the other hand, at subinhibitory concentrations, secondary metabolites are also described as signaling molecules within microbial communities or influencers of cellular differentiation (2–4).

One of the most intensely studied species of soil bacteria is *Bacillus subtilis* that serves as a laboratory model organism for biofilm formation and sporulation (5). *B. subtilis* is the type species of the *B. subtilis* species complex, containing the phylogenetically and phenetically homogeneous species *B. subtilis*, *Bacillus amyloliquefaciens*, *Bacillus licheniformis* and *Bacillus pumilus* (6). Several studies have shown, that members of the genus *Bacillus* produce various secondary metabolites of which many have bioactive properties (7, 8). These secondary metabolites, including polyketides, terpenes, siderophores and both ribosomal and non-ribosomal synthesized peptides are encoded by large biosynthetic gene clusters (BGCs) (9). While numerous natural products have been identified in the *B. subtilis* species complex, the diversity of secondary metabolite production in numerous isolates from the same niche has not been explored to understand their ecological functions.

Furthermore, it has been shown that *Bacillus* sp. have excellent biocontrol properties by promoting plant growth and reducing plant diseases caused by both plant pathogenic fungi and bacteria (10). These properties are mostly linked to their secondary metabolite profiles. One very potent chemical group of secondary metabolites are non-ribosomally produced lipopeptides which have various antimicrobial properties. *Bacillus* sp. produce different lipopeptide isoforms belonging to the families of surfactins, fengycins or iturins (11). A comparison of predicted BGCs from distinct *Bacillus* genomes assigned 11 BGCs to *B. subtilis* strains (12). Notably, a predicted BGC is not proof of the synthesis of the natural product. Gene silencing or absence of unidentified environmental triggers can be a reason for the lack of BGC expression (9).

In this study, we focus on *sfp*-dependent non-ribosomal peptides (NRPs) produced by recently isolated *B. subtilis* soil isolates. NRPs are synthesized by large enzyme-complexes, non-ribosomal peptide synthetases (NRPSs) (9). *B. subtilis* harbors three NRPS gene clusters (surfactin, plipastatin and bacillibactin) and one hybrid NRPS-PKS gene cluster (bacillaene). The phosphopantetheinyl transferase Sfp plays an important role in the NRP syntheses in *B. subtilis*, since it functions as an activator of the peptidyl carrier protein domains, converting it from the inactive *apo*-form to the active *holo*-form by transferring the 4-phosphopantetheine of Coenzyme A as prosthetic group to a conserved serine residue (13). The domesticated laboratory strain 168 contains an inactive *sfp* gene due to a frameshift mutation, causing the incapability of NRP production (13–15). Surfactin, encoded by the *srfAA-AD* gene cluster, is a well-studied biosurfactant and is involved in reducing surface tension needed for swarming and sliding motility (16, 17). Its cytolytic activity is mainly based on its surfactant activity, causing cell lysis due to penetration of bacterial lipid bilayer membranes and forming ion-conducting channels (9, 18–23). The powerful antifungal lipopeptide plipastatin, chemically very similar to fengycin, but distinct D-tyrosine position within the peptide backbone, is synthesized by the *ppsA*-*E* gene cluster. However, it was recently shown that *B. subtilis*, unlike *Bacillus velezensis* and *Bacillus amyloliquefaciens* mainly produces plipastatin and not fengycin (24). The detailed mode of action is not yet investigated but it is believed that it functions as an inhibitor of phospholipase A2, forming pores in the fungal membrane and causing morphological changes in fungal membrane and cell wall (9, 25, 26). Many studies have shown that plipastatin and fengycin are bioactive against diverse filamentous fungi (23, 27–32). Bacillaene, expressed from the *pksA-S* gene cluster, is a broad-spectrum antibiotic acting by inhibiting bacterial protein synthesis, but can also protect cells and spores from bacterial predators (33, 34). Bacillibactin, synthesized by the *dhbA*-*F* gene cluster, is a siderophore and transports iron from the environment to the cell (35). However, no studies have been published on its antimicrobial properties.

Most studies in the literature concentrate on single *Bacillus* sp. isolates, often discovered due to excellent antimicrobial properties. In this study, we focused on recently and systematically isolated *B. subtilis* environmental strains to get a more comprehensive overview of the chemodiversity within the species. We concentrated on differences in their antifungal properties, NRP production and genomic background of secondary metabolite arsenal. The antifungal properties of natural isolates and their respective NRP mutant derivatives were tested by antagonism assays with the three plant pathogenic fungi *Fusarium oxysporum*, *Fusarium graminearum* and *Botrytis cinerea*. *F. oxysporum* is a known plant pathogenic fungus causing *Fusarium* wilt in tomato, tobacco or banana plants amongst others (36). *F. graminearum* causes *Fusarium* head blight in different cereal crops (37). *B. cinerea* has a very wide variety of hosts, however, its main hosts are wine grapes and other fruits where it is causing the grey mold disease (38, 39). Using a library of *B. subtilis* isolates, we identified the NRPs that are responsible for inhibiting *Fusarium* sp. and *Botrytis cinerea*. Further, using fungal inhibition profiles, chemical detection of the NRPs, and detailed genomic analysis, we discovered that isolates originating from the same soil sample site possess distinct secondary metabolite production abilities suggesting chemical differentiation of *B. subtilis* in the environment.

## RESULTS

### Testing the antifungal potential of *B. subtilis* isolates

A small library of *B. subtilis* isolates has been established from various locations in Denmark and Germany (see Materials and Methods). To confirm that NRPs produced by these *B. subtilis* soil isolates have antifungal potential, we screened both wild type isolates and their *sfp* mutants (Fig. 1A) as well as their single NRP mutant derivatives *srfAC*, Δ*ppsC* and Δ*pksL* (Fig. 1B) against the three plant-pathogenic fungal strains *F. oxysporum*, *F. graminearum* and *B. cinerea*. The mutant screen allowed us to verify if a single NRP or a mixture of them is responsible for the bioactivity.

**FIG 1.**
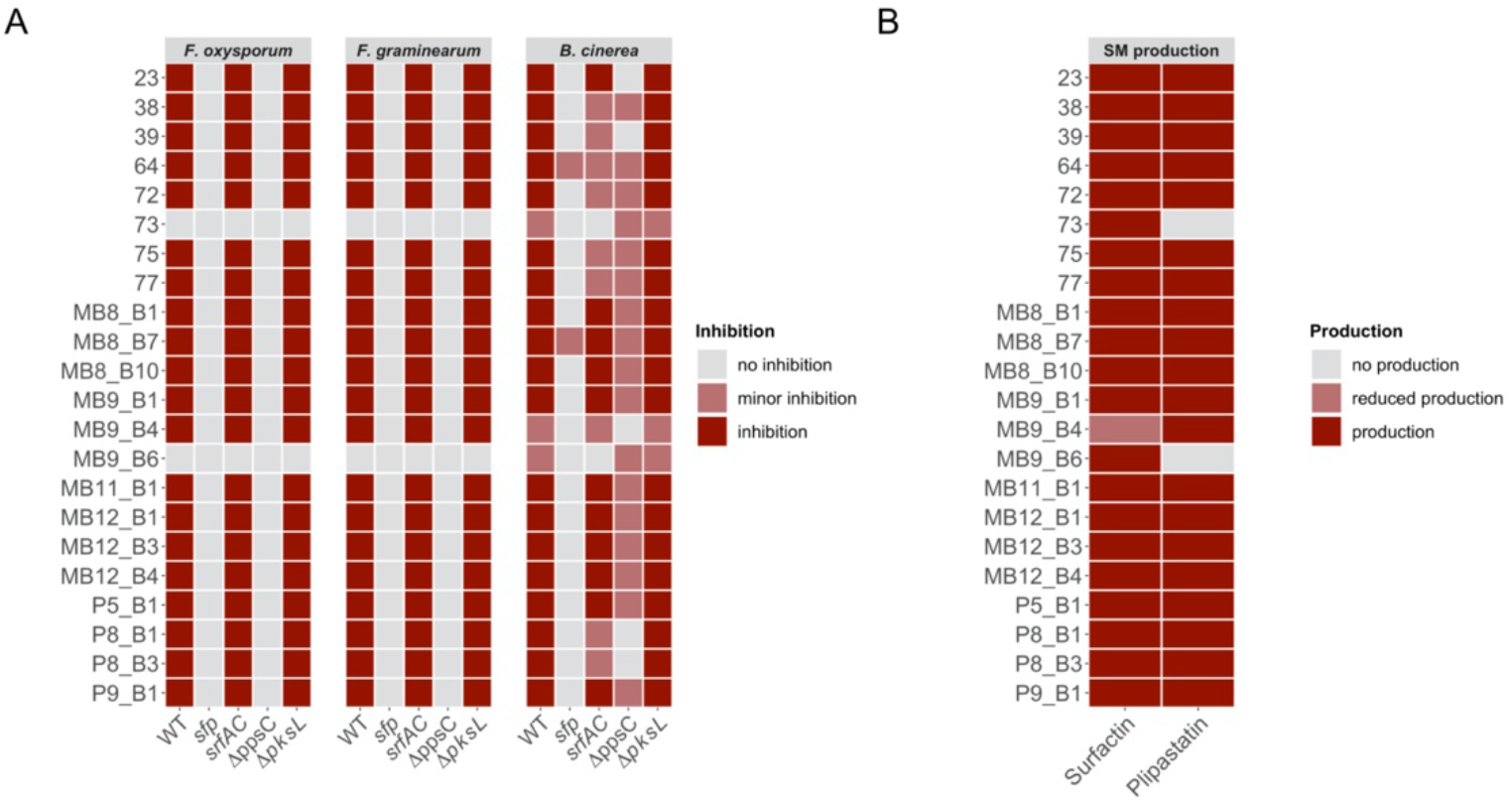
(A) Overview of qualitative evaluation of antagonisms assays assigned to inhibition, minor inhibition and no inhibition. (B) Overview of the detected non-ribosomal peptides surfactin and plipastatin in the extracts of wild type soil isolates by ESI–MS.

The qualitative assessment of antifungal potential from 22 tested isolates was classified into inhibition, minor inhibition and no inhibition by comparing mutant strains to their respectively wild types and wild types with each other (Fig. 2A, Fig. S2). The incidence of a distinct inhibition zone was defined as inhibition, while no inhibition refers to a total loss of inhibitory potential which appeared in surrounded or overgrown bacterial colonies by the fungus.

**FIG 2.**
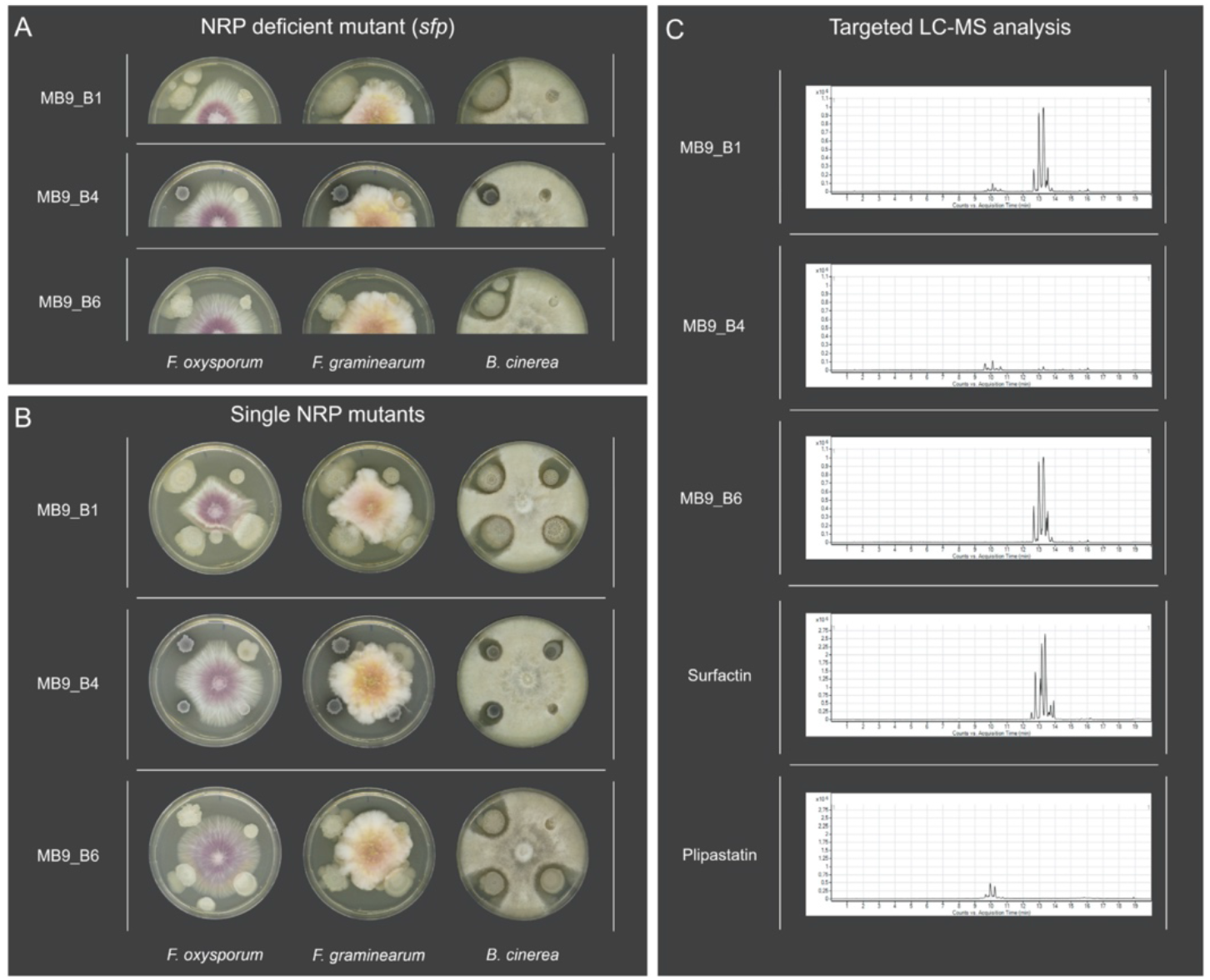
(A) Antagonism assays between the plant pathogenic fungi *Fusarium oxysporum*, *Fusarium graminearum* and *Botrytis cinerea*, and the *B. subtilis* soil isolates (left) as well as their NRP deficient *sfp* mutants (right). (B) Antagonism assays between the plant pathogenic fungi and *B. subtilis* soil isolates (upper left) as well as their single non-ribosomal peptide mutants *srfAC* (upper right, no surfactin), Δ*ppsC* (lower right, no plipastatin) and Δ*pksL* (lower left, no bacillaene). 5 μl of bacterial overnight culture and fungal spore suspension was spotted on the edges (bacteria) and in the center (fungi) of potato dextrose agar (PDA) plates. Strains were co-cultivated at 21-23°C for 6 days. (C) Extracted ion chromatograms (m/z 1000–1600) display various production levels of surfactin and plipastatin among *B. subtilis* soil isolates. The standard mixtures of plipastatin and surfactin are shown below.

To increase the possibility of differentiating between slight distinctions, we assigned strains exhibiting a reduced antagonism to the class minor inhibition. This observation differed between mutant derivatives and WTs. We specified minor inhibitions for WT, *srfAC*, Δ*ppsC* and Δ*pksL* strains when a thin layer of fungal hyphae was growing into the visible clearing zone towards the bacterial colony. In contrast, for *sfp* mutants, it described a not entirely loss of bioactivity (Fig. S1). Two isolates P5_B2 and P8_B2 were excluded from further antifungal screening since these strains were not naturally competent, and therefore we were unable to create NRP mutants. However, both wild types showed no inhibition of *Fusarium* sp. or *B. cinerea*.

20 of 22 tested wild types showed inhibition of *F. oxysporum* and *F. graminearum*, whereas their *sfp* mutants showed no growth inhibition. Exceptions were the strains 73 and MB9_B6 which showed no antagonistic effects against *Fusarium*. For all 20 bioactive strains, only their Δ*ppsC* mutants, incapable of producing plipastatin, lost the bioactivity against both *Fusarium* species comparable to the *sfp* mutants.

The screening revealed that *B. cinerea* is not as sensitive to a single compound as observed in the case of *Fusarium*. All tested WT strains inhibited *B. cinerea*, merely three strains showed minor inhibition. Similar to the *Fusarium* antagonism, strains incapable of producing NRPs (*sfp* mutant) lost their abilities to hinder *B. cinerea* growth. Only two of the *sfp* mutants showed remaining antagonistic effect albeit reduced compared to their respective WTs. The inactivation of either plipastatin or surfactin production caused different screening results depending on the specific soil isolate. Five of the strains showed minor inhibition or unaffected inhibition when surfactin production was disrupted, but a total loss of inhibition when the plipastatin BGC was absent in a given strain. Another group of five strains displayed only minor inhibitions independent of the lack of BGCs for surfactin or plipastatin. The majority of tested strains (i.e. 10 isolates), were not affected by the inactivation of surfactin production, however, the lack of plipastatin BGC caused minor inhibition. As an exception, two strains exhibited a total loss of bioactivity against *B. cinerea* without the surfactin BGC but still minor inhibition when plipastatin gene cluster was lacking. None of the bacillaene (Δ*pksL*) mutants displayed any changes compared to their WTs.

Among all strains, three strains stand out in the antifungal screen. Strains 73 and MB9_B6 lacked inhibition of *F. oxysporum* and *F. graminearum* and displayed reduced inhibition of *B. cinerea*. Both their *sfp* and *srfAC* derivatives showed a complete loss of bioactivity against *B. cinerea*. In contrast to these strains, isolate MB9_B4 showed no antifungal inhibition when plipastatin BGC was disrupted.

### Chemical characterization of *B. subtilis* isolates and their mutant derivatives

Screening of antifungal activities revealed potential differences in surfactin and plipastatin production among the isolates. Therefore, to compare the production of these NRPs among the soil isolates, a targeted LC-MS analysis was performed targeting compounds with m/z values between 1000 and 1600 (40, 41). Interestingly, even co-isolated strains showed varying NRP production (Fig. 1C). The qualitative analysis disclosed that the majority of strains produced both surfactin and plipastatin (Fig. 2B, Fig. S3), while the three peculiar strains from the antifungal screening have a distinct natural product profile. Plipastatin was not detectable in the extracts from strains 73 and MB9_B6. Absence of plipastatin production by these isolates correlates with lack of antagonism against the *Fusarium* species and reduced *B. cinerea* inhibition compared to other isolates. Additionally, strain MB9_B4 with strongly lowered surfactin level displayed a reduced inhibition of *B. cinerea*. Importantly, its plipastatin mutant exhibited a total loss of antagonism. The results demonstrate that if wild types do not produce plipastatin or surfactin and production of the counterpart NRP is genetically hindered, strains lose the bioactivity against *B. cinerea*.

### Synergism between plipastatin and surfactin for *B. cinerea* inhibition

The deletion mutant screen in combination with chemical profiling suggested that the presence of either plipastatin or surfactin is sufficient for inhibiting the growth of *B. cinerea*. To test this hypothesis, both BGCs were disrupted in strains that were found to be capable of producing both compounds in chemical profiling. However, all three tested *srfAC*-Δ*ppsC* double-mutants maintained bioactivity against the fungus (Fig. S4A), even though targeted LC-MS analysis of two of these tested strains confirmed the lack of both lipopeptides (Fig. S4C). These data indicate that either the bioactivity is not only caused by surfactin and plipastatin for some of the strains or intermediates secreted in these mutants preserve the bioactive properties that hamper *B. cinerea* growth.

### Prediction of biosynthetic gene cluster potential of the isolates from their genome sequences

Based on the antifungal screening results and the origin of the isolate, 13 *B. subtilis* strains were selected for genome sequencing, in addition to one *B. licheniformis* strain used as an outlier (42). The genomes were analyzed with antiSMASH (43) to obtain an overview of all predicted BGCs (Fig. 3) and their similarity to known clusters. Importantly, these predictions highlight the genomic potential, but not the actual production of secondary metabolites. Additionally, the whole BGCs and not solely the core genes are compared to gene clusters of appropriate reference strains.

**FIG 3.**
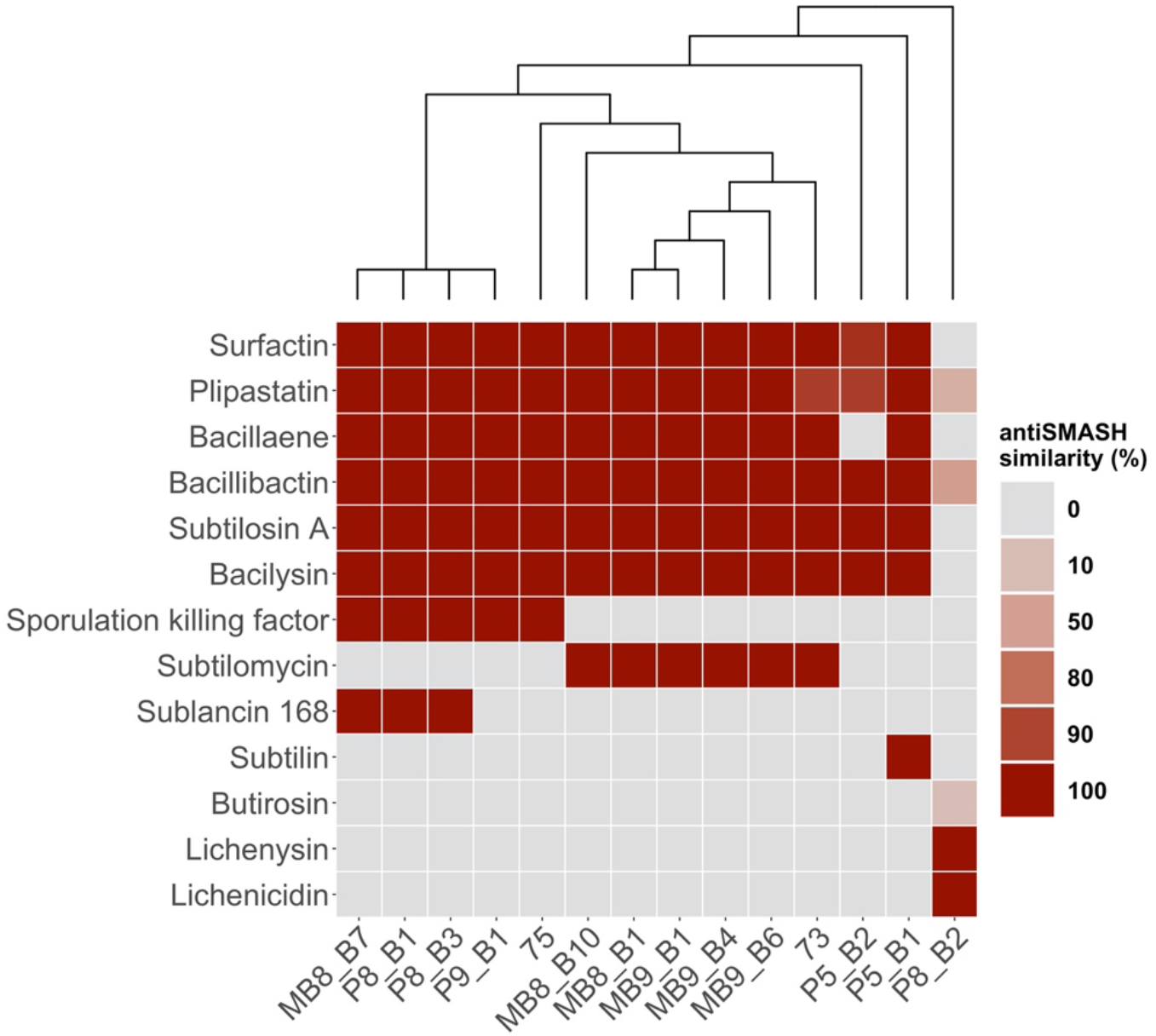
Overview of predicted biosynthetic gene clusters (BGCs) by antiSMASH of 13 *B. subtilis* and one *B. licheniformis* (right) soil isolates. Color-code visualizes the similarity of BGCs to a reference BGC, whereby the grey color (0 %) indicates their absence. The cladogram is based on a core gene alignment by the pan-genome pipeline Roary.

The BGCs for the *sfp*-dependent non-ribosomal peptides surfactin, plipastatin, bacillaene and bacillibactin were predicted in all *B. subtilis* isolates except for the bacillaene BGC absent in isolate P5_B2. The surfactin gene cluster shows for the majority of strains a similarity of 100 % compared to the reference, while only isolate P5_B2 exhibit a lower similarity due to minor differences in genes of the gene cluster. Likewise, for the plipastatin gene cluster, the greater number of *B. subtilis* strains showed a similarity of 100 %, except strains 73 and P5_B2 which both displayed absent genes compared to the reference gene cluster. Bacillibactin, subtilosin A and bacilysin are present in all *B. subtilis* strains with a similarity of 100 %. The sporulation killing factor is present in five strains, but these are lacking the gene cluster for subtilomycin production. On the contrary, six strains are predicted to code for subtilomycin synthesis and are vice versa missing the genes of the sporulation killing factor. Phelan *et al.* observed, that the subtilomycin gene cluster of a marine isolate is present in the genomic locus of the sporulation killing factor gene cluster (44). Finally, neither sporulation killing factor nor subtilomycin synthesis are predicted to be present in strains P5_B1 and P5_B2, however, subtilisin genes seem to be present in P5_B1. Strain P5_B2 represents an outlier among the *B. subtilis* isolates, since it possesses only the BGCs for the core secondary metabolites of *B. subtilis*, except bacillaene, but none of the accessory BGCs differentially present in the others. In line with the inhibition data, *B. licheniformis* P8_B2 has a deviating profile of BGCs. Three gene clusters show similarity to plipastatin, bacillibactin and butirosin with 30 %, 53 % and 7 %, respectively. Additionally, the BGCs for the species-specific secondary metabolites, lichenysin and lichenicidin were predicted in P8_B2 with 100 % similarity.

### Detailed comparison of BGC structures explains lack of NRP production

Differences in both the antifungal potential and plipastatin or surfactin production based on the targeted LC-MS analysis combined with the BGC prediction with antiSMASH led us to concentrate more on the non-producer or predicted non-producer strains MB9_B4, 73, MB9_B6 and P5_B2. To understand why these strains show these characteristics, the core genes of surfactin, plipastatin and bacillaene were analyzed in their presence and absence and compared to the BGCs of co-isolated producer strains (Fig. 4).

**FIG 4.**
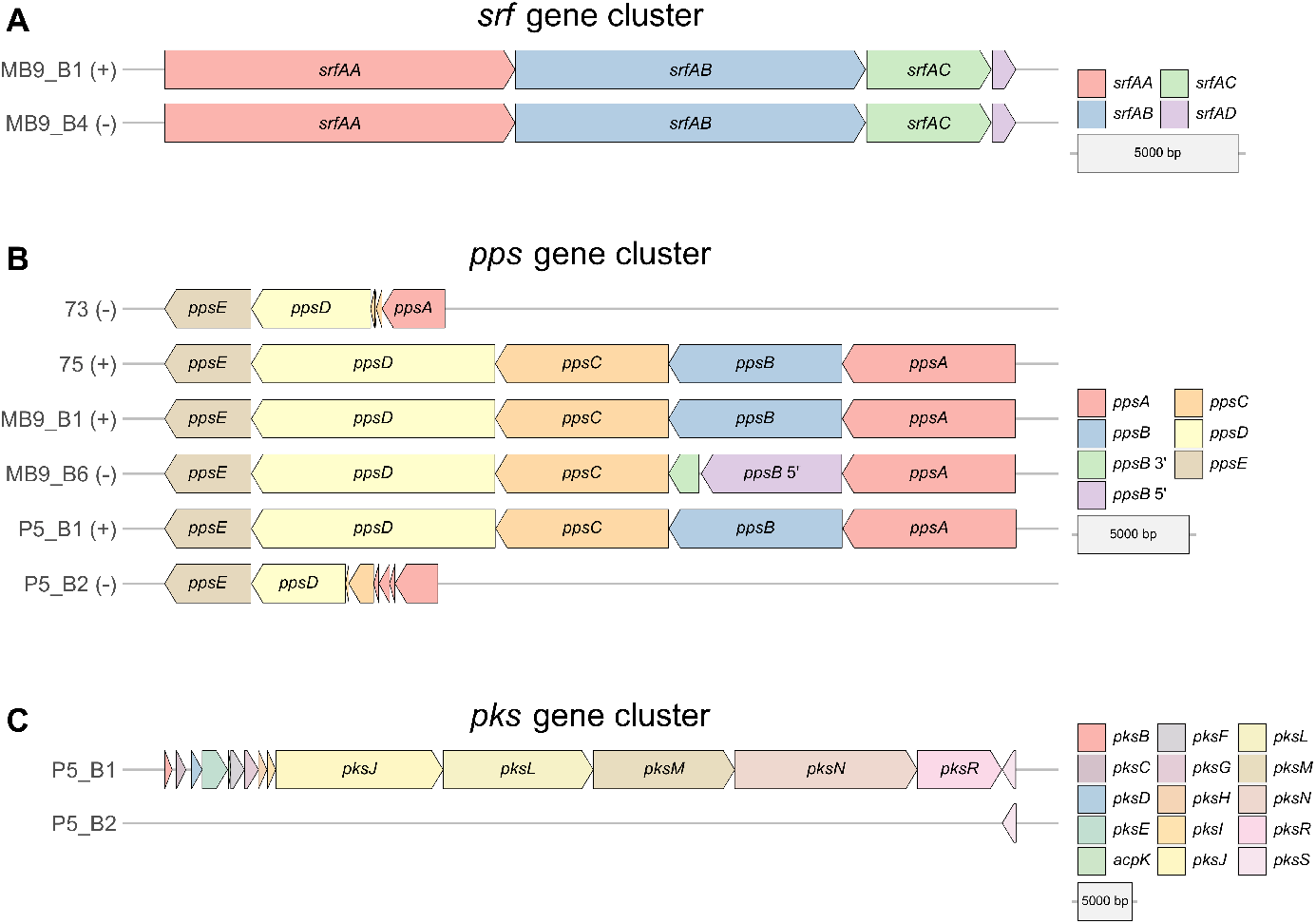
Comparison of core genes of the biosynthetic gene clusters surfactin (A) and plipastatin (B) from co-isolated *B. subtilis* producer (+) and non-producer (-) strains. (C) Comparison of core genes of the biosynthetic gene clusters of bacillaene from two co-isolated *B. subtilis* strains.

MB9_B4 showed a hampered surfactin production, even though all core genes of the surfactin BGC are present equally to the co-isolated producer strain MB9_B1 (Fig. 4A). We further analyzed genes involved in the regulation of surfactin BGC transcription. The comparison of *comA* of all 13 *B. subtilis* strains revealed six mutated regions, however, five of them were silent mutations. Interestingly, the point mutation at nucleotide position 3 is unique for MB9_B4 and causing an alteration in the translation initiating methionine (Fig. S5B). Consequently, the coding region of *comA* is reduced by 13 amino acids. It was shown, that the response regulator ComA triggers directly the transcription of the *srfA* operon by binding to its promoter region (45–47). The conserved residues at the amino acid position 8 and 9 that are reported to be one out of three targets for ComP-catalyzed phosphorylation are not translated due to the mutation in MB9_B4 (48). The results led us to assume, that surfactin production of MB9_B4 might be hampered due to altered regulatory processes.

Plipastatin was not detectable in the strains 73, MB9_B6 and P5_B2 and all were incapable of inhibiting the tested *Fusarium* strains. Analyses of the *pps* gene cluster from strains 73 and P5_B2 revealed the presence of only smaller fragments of the genes *ppsA*, *ppsC* and *ppsD*, a complete absence of *ppsB*, but a present *ppsE* gene (Fig. 4B). Interestingly, MB9_B6 is harboring all five *pps* core genes, however, the translation of the *ppsB* gene is interrupted by a point-nonsense mutation G→A which causes an amino acid change from tryptophan to a termination codon (Fig. S5A). The resulting non-translated region of 41 amino acids results in dysfunction of the carrier and epimerization regions of the second domain. Therefore, the plipastatin production is most likely inactive due to either missing core genes or a disrupted *ppsB* gene.

In this study, we have not measured the production of bacillaene, nevertheless, strain P5_B2 is missing all core genes of the *pks* gene cluster except *pksS* (Fig. 4C). This genomic background strongly supports the assumption, that P5_B2 is incapable of producing bacillaene.

We conclude that differences in the synthesis of surfactin, plipastatin and bacillaene are caused due to regulatory processes, a disrupted core gene caused by a point-nonsense mutation or a loss of several core genes, highlighting the diversity of NRP production in soil isolates of *B. subtilis*.

### Intraspecies interactions of soil isolates

In addition to the antifungal activities, strains isolated from the same sampling site were co-cultivated to determine if they inhibit one another (Fig. 5A). This allowed examination of co-isolates from 5 different sample sites. These results were compared to their BGC profiles predicted by antiSMASH (Fig. 5B). Importantly, none of the strains showed self-inhibition, except MB8_B10 that displayed a clear inhibition zone. Strain 73 inhibited strain 75, but not vice versa. The BGC of the lantibiotic subtilomycin is predicted for strain 73 but absent in strain 75. Both MB8_B1 and MB8_B10 have a predicted subtilomycin BGC and inhibited MB8_B7. Furthermore, these strains showed only minor inhibitions to each other. MB8_B7 is inhibiting both MB8_B1 and MB8_B10, which can be traced back to the predicted BGC of sublancin. MB9 strains showed one common BGC, subtilomycin and in line with this, no inhibition was detectable during the antagonism screens. P5_B1 harboring a predicted subtilin BGC inhibited P5_B2, which has none of the targeted BGCs predicted. Nevertheless, P5_B2 showed still a reduced inhibition of P5_B1. In line with antiSMASH predicting the presence of sublancin BGC in both P8_B1 and P8_B3, these isolates inhibited their co-inhabitant P8_B2. This *B. licheniformis* strain, P8_B2 had none of the targeted BGCs predicted, but the *B. licheniformis*-specific BGCs of butirosin, lichenysin and lichenicidin. However, P8_B2 was inhibiting P8_B1 and showed minor inhibition of P8_B3. Based on the screening results and predicted BGCs, we can assume that strains having the same BGCs were not inhibiting each other or showed only minor inhibition. Moreover, strains with different BGCs had a variable inhibitory effect on each other, possibly due to the lack of resistance genes for the specific secondary metabolite.

**FIG 5.**
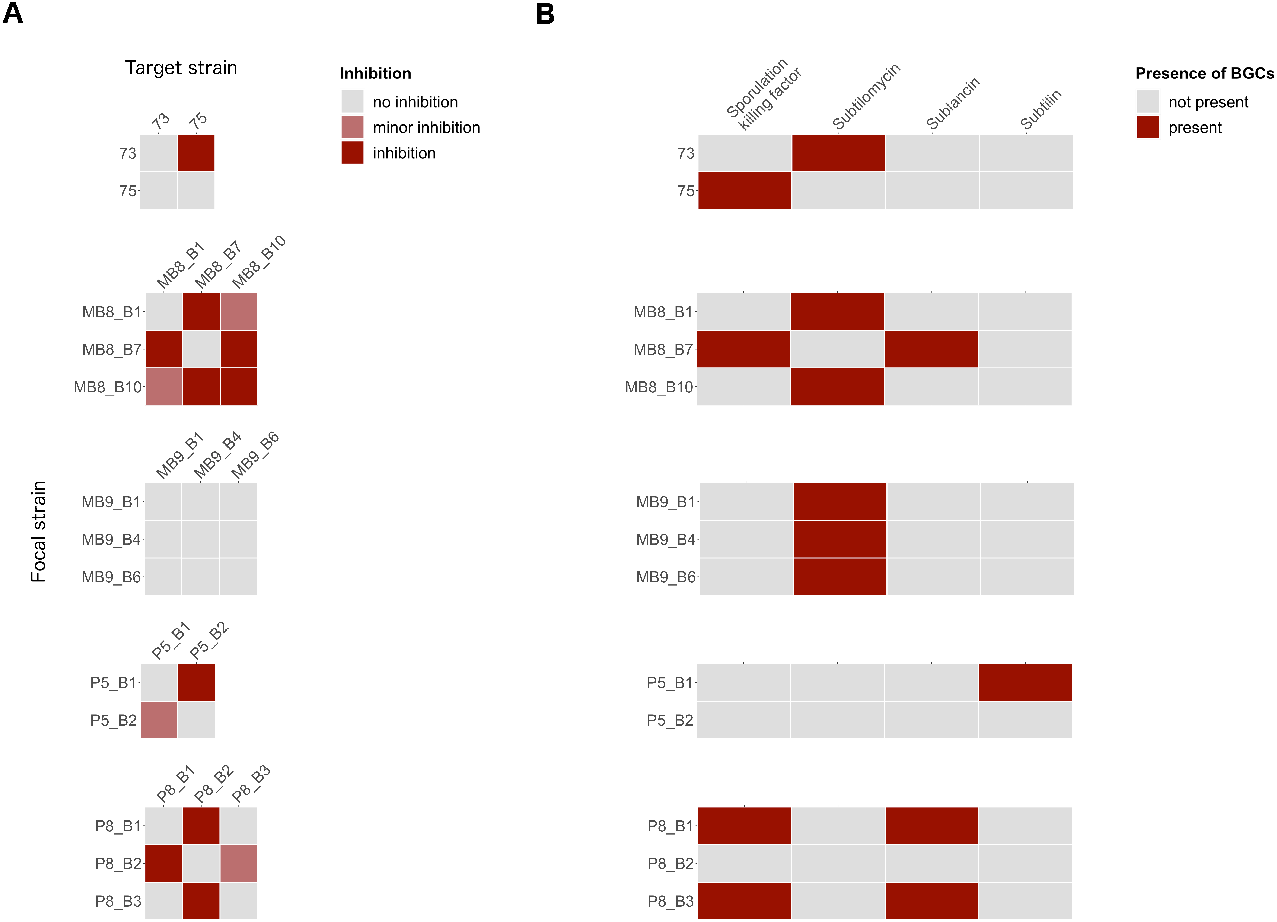
(A) Overview of intra-species interactions of co-isolated *B. subtilis* strains. Target strains were embedded in 1 % LB agar and 8 μl of focal strains were spotted on top. Plates were incubated at 37°C for 24 h. (B) Overview of predicted accessory BGCs by antiSMASH.

## DISCUSSION

Undomestica isolates of *B. subtilis* produce a wide range of different secondary metabolites, defining their biocontrol properties. The produced secondary metabolites affect fungal and bacterial growth and differentiation, and possibly other macroorganisms. Our study provides an overview of the antifungal properties and secondary metabolite profiles of recently and partly co-isolated environmental strains of *B. subtilis*.

Our screening results revealed that antifungal properties vary among *B. subtilis* soil isolates and interestingly also among co-isolated strains from the same soil sample. We demonstrated that only plipastatin is necessary to inhibit the growth of *F. oxysporum* and *F. graminearum*. Plipastatin is known as a powerful fungicide by inhibiting the phospholipase A2 and forming pores in fungal membranes (9, 25, 26). On the contrary, the anti-*Botrytis* potential of *B. subtilis* is linked to multiple *sfp*-dependent NRPs, both surfactin and plipastatin contribute to full fungal inhibition. Disrupting the BGC responsible for the production of surfactin or plipastatin in the strain that produces only one of these NRPs eliminated the strains’ anti-*Botrytis* activity, while the mutating of both BGCs in a strain that originally produces both surfactin and plipastatin still maintains slight activity against *B. cinerea*, despite a clear *sfp*-dependent inhibition. In the case of both BGC mutants, only one gene in the respective BGC was knocked out, thus potential intermediates might be produced that affect the fungal growth. Further purification of bioactive natural products will be necessary to reveal the chemical nature of the inhibitory molecule. Plausibly, these strains might additionally produce cell wall degrading enzymes or volatiles which could contribute to the observed remaining antifungal activity. Nevertheless, production of plipastatin or a combination of both plipastatin and surfactin are important in *B. subtilis* to suppress *B. cinerea* growth.

Genetic differentiation and loss of secondary metabolite production is a rapid process observed previously with laboratory strains of *B. subtilis*. The most widely used *B. subtilis* model strains (e.g. 168 and PY79) have been observed to rapidly lose their ability to produce NRPs during domestication due to a mutation in the *sfp* gene (49). The lack of surfactin production reduced swarming, therefore easy cultivation on agar media probably influenced domestication of this species.

Interestingly, hampered surfactin production was noticed in isolate MB9_B4. We hypothesize that the reduction of surfactin production might be due to altered gene regulation by a mutation in the *comA* gene. However, it is unclear to what extend the ComA protein level is affected due to mutation in the translation initiating methionine and whether the potentially altered level of this transcription factor in strain MB9_B4 mitigates sufficient binding to the promoter region of *srf* to activate its transcription (50). Repair of the *comA* mutation could potentially corroborate if this single mutation is causing the reduced surfactin production in MB9_B4. Notably, the *srf* operon also codes for the anti-adaptor protein ComS, which is required for competence development (51). The reduced expression of *srf* gene cluster would also attenuate the co-transcription of *comS*, therefore causing diminished competence. However, we found no evidence that competence is affected in MB9_B4 compared to other isolates.

Besides reduced surfactin production, plipastatin was not detected in the extracts of three isolates: 73, MB9_B6 and P5_B2. A point-nonsense mutation could be identified in the *ppsB* gene of strain MB9_B6, which possibly hinders the assembly of a functional plipastatin producing complex. In contrast, the core genes *ppsA-C* and a part of *ppsD* are absent in strains 73 and P5_B2 resulting in lack of plipastatin production. Intriguingly, these two strains that were isolated from soil samples in Germany and Denmark, carry very similar deletions in the pps gene cluster. Finally, strain P5_B2 lost almost the complete *pks* gene cluster.

These intriguing examples of partial BGCs in environmental *B. subtilis* strains highlight the possibility for secondary metabolite production loss in nature. Moreover, these observations suggest that either the selection pressure is not strong enough to maintain the production of these specialized metabolites in the particular niches or that secondary metabolites can be shared as common goods in bacterial populations, therefore these derivatives can act as cheaters. Besides, the appearance of producer and non-producer strains in a bacterial population can be also seen as a division of labor, as suggested to be present in the *Streptomyces* genus (52). It remains to be examined whether the derivatives with mutated secondary metabolite production has altered fitness when growing in soil. However, the natural role of most secondary metabolites is ambiguous; thus, gene loss in these BGCs in nature might be not as detrimental as expected from *in vitro* laboratory observations.

## MATERIAL AND METHODS

### Strains, media and chemicals

All strains that were used in this study or that were used solely as gDNA donors for transformation are listed in Table S1. For routine growth, bacterial cells were cultured in Lysogeny Broth medium (LB-Lennox, Carl Roth, Germany; 10 g l^−1^ tryptone, 5 g l^−1^ yeast extract and 5 g l^−1^ NaCl) supplemented with 1.5 % Bacto agar if required. When necessary, antibiotics were used at the following concentrations: MLS (1 μg mL^−1^ erythromycin, 25 μg mL^−1^ lincomycin); spectinomycin (100 μg mL^−1^); chloramphenicol (5 μg mL^−1^), tetracycline (10 μg mL^−1^), erythromycin (5 μg mL^−1^) and ampicillin (100 μg mL^−1^). Soil isolates were obtained from 11 sampling sites in Germany and Denmark (see Table S1 for coordinates) by selecting for spore formers in the soil. Soil samples were mixed with 0.9 % saline solution, vortexed on a rotary shaker for 2 min, incubated at 80 °C for 25 min and serially diluted on LB medium solidified with 1.5 % agar (53). Highly structured colonies were targeted and isolation of *B. subtilis* strains was confirmed using 16S sequencing followed by whole-genome sequencing of 13 selected strains (42) and one additional isolate identified as *B. licheniformis*.

*F. oxysporum*, *F. graminearum* and *B. cinerea* were revived on Potato Dextrose Agar (PDA, BD, USA, potato infusion 4 g l^−1^, glucose 20 g l^−1^, agar 15 g l^−1^) supplemented with 0.5 g l^−1^ CuSO_4_ and 0.5 g l−1 ZnSO_4_ to harvest spores.

### *B. subtilis* mutant strain construction

Strains 23, 38, 39, 64, 72, 73, 75 and 77 were isolated specifically by labelling with constitutively expressed *gfp* from P_hyperspank_ using phyGFP plasmid that integrates into the *amyE* locus (54). Mutant strains were obtained using natural competence (55) by transforming genomic DNA and selecting for antibiotic resistance, followed by verifying the mutation by PCR. Mutants were constructed by transforming gDNA of the following strains: *sfp* mutants from DS3337 (56), *srfAC* mutants from DS1122 (57), Δ*pksL* mutants from DS4085 (34) and Δ*ppsC* mutants from DS4114 (34). The *srfAC*-Δ*ppsC* double mutants were obtained by transforming gDNA from DS4114 (34) into the respective *srfAC* mutants.

### Antagonism assays between plant pathogenic fungi and *B. subtilis* soil isolates

Spores of fungal cultures grown at 21-23°C for 5–7 days on sporulation medium were harvested using 10 ml saline tween solution (8 g l^−1^ NaCl and 0.05 ml l^−1^ Tween 80) and filtered through Miracloth (Millipore; Billerica, USA) following the protocol described from Benoit *et al*. (58). The spore solution was centrifuged 5000 rpm for 10 min, resuspended in saline tween solution and stored at 4°C until use. 5 μl of bacterial overnight cultures and fungal spore suspension were spotted on the edge (bacteria) and in the center (fungus) of Potato Extract Glucose Agar (PDA) plates (Carl Roth, Germany; potato infusion 4 g l^−1^, glucose 20 g l^−1^, agar 15 g l^−1^, pH value 5.2 ± 0.2). Plates were cultivated at 21-23°C for 6 days and antagonistic observations were qualitatively documented.

### Extraction of secondary metabolites

Bacterial strains were cultured on PDA plates and incubated at 30°C for 3 days. A 6 mm-diameter size agar plug of the bacterial culture was transferred to a 2 mL size Eppendorf tube and extracted with 1 mL organic solvent (2-propanol:EtOAc (1:3, v/v) containing 1 % formic acid). The tubes were then sonicated for 60 min. The solutions were transferred to new tubes, evaporated under N_2_, and redissolved in 300 μL of MeOH before further sonication for 15 min, followed by 3 min of centrifugation (13,400 rpm). After centrifugation, the supernatants were transferred to clean HPLC vials and subjected to ultrahigh-performance liquid chromatography-high resolution mass spectrometry (UHPLC-HRMS) analysis.

### UHPLC-HRMS analysis

A volume of 1 μL extract was subjected to UHPLC-HRMS analysis. UHPLC-HRMS was performed on an Agilent Infinity 1290 UHPLC system fitted with a diode array detector. Liquid chromatography was run on an Agilent Poroshell 120 phenyl-hexyl column (2.1 × 250 mm, 2.7 μm) at 60°C with MeCN–H_2_O gradient, both containing 20 mM formic acid. A linear gradient of 10 % MeCN/H_2_O to 100 % MeCN over 15 min was initially employed, followed by an isocratic condition of 100 % MeCN for 2 min before returning to starting conditions of 10 % MeCN/H_2_O for 3 min, all at a flow rate of 0.35 mL/min. An Agilent 6545 QTOF MS equipped with an Agilent Dual Jet Stream electrospray ion (ESI) source was used for MS detection in positive ionization. The MS detection was performed with a drying gas temperature of 250°C, drying gas flow of 8 L/min, sheath gas temperature of 300°C, and sheath gas flow of 12 L/min. The capillary voltage was set to 4000 V and nozzle voltage to 500 V. MS data were processed and analyzed using Agilent MassHunter Qualitative Analysis B.07.00.

### Intraspecies interactions

Bacterial o/n-cultures were adjusted to an OD of 2. LB plates were prepared with the agar overlay technique: 10 mL LB medium containing 1.5 % agar functioned as a bottom layer and was overlaid by 10 mL LB medium containing 1 % agar that was pre-inoculated with the target strain in a 1:200 dilution. 8 μL of the focal strain was spotted on the 25 min pre-dried double-layer plates and incubated at 37°C for 24 h. Interactions were evaluated by checking appeared clearing zones between the focal colonies and bacterial lawns of the target strains.

### Bioinformatic analysis

The genomes of 14 selected soil isolates (13 *B. subtilis*, 1 *B. licheniformis*) published in Kiesewalter *et al.*, 2020 (42) were submitted to antiSMASH 5.0 to analyze the differences in their gene cluster predictions (43). The pan-genome pipeline Roary was applied to the Prokka annotations of the *B. subtilis* genomes to construct a pan-genome of genes having 95% similarity in 99% of the isolates (59, 60). The presence and absence gene list generated by Roary was used for comparisons between selected BGCs. Single gene comparisons were conducted by aligning both the nucleotide sequences and, with seqKit (61), translated amino acid sequences with MUSCLE (62) and inspecting them in Jalview 2 (63). All gene clusters or single genes were visualized in R by using the publicly available ggplot2 extensions gggenes, ggseqlogo and ggmsa (64–67). A phylogenetic tree was calculated with FastTree 2 using the core gene alignment by Roary and visualized in R with the publicly available ggplot2 extension ggtree (68, 69).

## ACKNOWLEDGEMENT

This project was supported by the Danish National Research Foundation (DNRF137) for the Center for Microbial Secondary Metabolites. T.W. and T.S.J. are funded by grants of the Novo Nordisk Foundation (NNF10CC1016517, NNF16OC0021746).

H.T.K. and Á.T.K. designed research; H.T.K, C.N.L.-A., M.W., D.S. performed research; H.T.K, C.N.L.-A., M.W., M.L.S, T.S.J. analyzed the data; G.M., T.O.L., V.S.C, T.W. contributed analytical tools; H.T.K. and Á.T.K. wrote the manuscript; all authors approved the manuscript.

## REFERENCES

1. Foster KR, Bell T. 2012. Competition, not cooperation, dominates interactions among culturable microbial species. Curr Biol 22:1845–1850.

2. Romero D, Traxler MF, López D, Kolter R. 2011. Antibiotics as signal molecules. Chem Rev 111:5492–5505.

3. Linares JF, Gustafsson I, Baquero F, Martinez JL. 2006. Antibiotics as intermicrobiol signaling agents instead of weapons. Proc Natl Acad Sci U S A 103:19484–19489.

4. Straight PD, Willey JM, Kolter R. 2006. Interactions between *Streptomyces coelicolor* and *Bacillus subtilis*: Role of surfactants in raising aerial structures. J Bacteriol 188:4918–4925.

5. Kovács ÁT. 2019. Bacillus subtilis. Trends Microbiol 27:724–725.

6. Fritze D. 2004. Taxonomy of the genus *Bacillus* and related genera: The aerobic endospore-forming bacteria. Phytopathology 94:1245–1248.

7. Stein T. 2005. *Bacillus subtilis* antibiotics: Structures, syntheses and specific functions. Mol Microbiol 56:845–857.

8. Kaspar F, Neubauer P, Gimpel M. 2019. Bioactive secondary metabolites from *Bacillus subtilis*: A comprehensive review. J Nat Prod 82:2038–2053.

9. Harwood CR, Mouillon JM, Pohl S, Arnau J. 2018. Secondary metabolite production and the safety of industrially important members of the *Bacillus subtilis* group. FEMS Microbiol Rev 42:721–738.

10. Fan B, Blom J, Klenk HP, Borriss R. 2017. *Bacillus amyloliquefaciens*, *Bacillus velezensis*, and *Bacillus siamensis* form an “operational group *B. amyloliquefaciens*” within the *B. subtilis* species complex. Front Microbiol 8:22.

11. Ongena M, Jacques P. 2008. *Bacillus* lipopeptides: versatile weapons for plant disease biocontrol. Trends Microbiol 16:115–125.

12. Grubbs KJ, Bleich RM, Santa Maria KC, Allen SE, Farag S, Shank EA, Bowers AA. 2017. Large-scale bioinformatics analysis of *Bacillus* genomes uncovers conserved roles of natural products in bacterial physiology. mSystems 2:e00040–17.

13. Quadri LEN, Weinreb PH, Lei M, Nakano MM, Zuber P, Walsh CT. 1998. Characterization of Sfp, a *Bacillus subtilis* phosphopantetheinyl transferase for peptidyl carder protein domains in peptide synthetases. Biochemistry 37:1585–1595.

14. Tsuge K, Ano T, Hirai M, Nakamura Y, Shoda M. 1999. The genes *degQ*, *pps*, and *lpa-8* (*sfp*) are responsible for conversion of *Bacillus subtilis* 168 to plipastatin production. Antimicrob Agents Chemother 43:2183–2192.

15. Nakano MM, Corbell N, Besson J, Zuber P. 1992. Isolation and characterization of *sfp*: a gene that functions in the production of the lipopeptide biosurfactant, surfactin, in *Bacillus subtilis*. MGG Mol Gen Genet 232:313–321.

16. Kearns DB, Losick R. 2003. Swarming motility in undomesticated *Bacillus subtilis*. Mol Microbiol 49:581–590.

17. Grau RR, De Oña P, Kunert M, Leñini C, Gallegos-Monterrosa R, Mhatre E, Vileta D, Donato V, Hölscher T, Boland W, Kuipers OP, Kovács ÁT. 2015. A duo of potassium-responsive histidine kinases govern the multicellular destiny of *Bacillus subtilis*. MBio 6:e00581–15.

18. Deleu M, Bouffioux O, Razafindralambo H, Paquot M, Hbid C, Thonart P, Jacques P, Brasseur R. 2003. Interaction of surfactin with membranes: A computational approach. Langmuir 19:3377–3385.

19. Sheppard JD, Jumarie C, Cooper DG, Laprade R. 1991. Ionic channels induced by surfactin in planar lipid bilayer membranes. BBA - Biomembr 1064:13–23.

20. Heerklotz H, Seelig J. 2001. Detergent-like action of the antibiotic peptide surfactin on lipid membranes. Biophys J 81:1547–1554.

21. Heerklotz H, Wieprecht T, Seelig J. 2004. Membrane perturbation by the lipopeptide surfactin and detergents as studied by deuterium NMR. J Phys Chem B 108:4909–4915.

22. Heerklotz H, Seelig J. 2007. Leakage and lysis of lipid membranes induced by the lipopeptide surfactin. Eur Biophys J 36:305–314.

23. Gao L, Han J, Liu H, Qu X, Lu Z, Bie X. 2017. Plipastatin and surfactin coproduction by *Bacillus subtilis* pB2-L and their effects on microorganisms. Antonie van Leeuwenhoek, Int J Gen Mol Microbiol 110:1007–1018.

24. Hussein W. 2019. Fengycin or plipastatin? A confusing question in Bacilli. Biotechnologia 100:47–55.

25. Umezawa H, Aoyagi T, Nishikiori T, Yamagishi Y, Okuyama A, Hamada M, Takeuchi T. 1986. Plipastatins: New inhibitors of phospholipase A2, produced by *Bacillus cereus* BMG302-fF67. J Antibiot (Tokyo) 39:737–744.

26. Deleu M, Paquot M, Nylander T. 2005. Fengycin interaction with lipid monolayers at the air-aqueous interface - Implications for the effect of fengycin on biological membranes. J Colloid Interface Sci 283:358–365.

27. Romero D, De Vicente A, Rakotoaly RH, Dufour SE, Veening JW, Arrebola E, Cazorla FM, Kuipers OP, Paquot M, Pérez-García A. 2007. The iturin and fengycin families of lipopeptides are key factors in antagonism of *Bacillus subtilis* toward *Podosphaera fusca*. Mol Plant-Microbe Interact 20:430–440.

28. Alvarez F, Castro M, Príncipe A, Borioli G, Fischer S, Mori G, Jofré E. 2012. The plant-associated *Bacillus amyloliquefaciens* strains MEP218 and ARP23 capable of producing the cyclic lipopeptides iturin or surfactin and fengycin are effective in biocontrol of sclerotinia stem rot disease. J Appl Microbiol 112:159–174.

29. Falardeau J, Wise C, Novitsky L, Avis TJ. 2013. Ecological and mechanistic insights into the direct and indirect antimicrobial properties of *Bacillus subtilis* lipopeptides on plant pathogens. J Chem Ecol 39:869–878.

30. Roy A, Mahata D, Paul D, Korpole S, Franco OL, Mandal SM. 2013. Purification, biochemical characterization and self-assembled structure of a fengycin-like antifungal peptide from *Bacillus thuringiensis* strain SM. Front Microbiol 4:332.

31. Tang Q, Bie X, Lu Z, Lv F, Tao Y, Qu X. 2014. Effects of fengycin from *Bacillus subtilis* fmbJ on apoptosis and necrosis in *Rhizopus stolonifer*. J Microbiol 52:675–680.

32. Zhang L, Sun C. 2018. Fengycins, cyclic lipopeptides from marine *Bacillus subtilis* strains, kill the plant-pathogenic fungus *Magnaporthe grisea* by inducing reactive oxygen species production and chromatin condensation. Appl Environ Microbiol 84:e00445–18.

33. Patel P, Huang S, Fisher S, Pirnik D, Aklonis C, Dean L, Meyers E, Fernandes P, Mayerl F. 1995. Bacillaene, a novel inhibitor of procaryotic protein synthesis produced by *Bacillus subtilis*: Production, taxonomy, isolation, physico-chemical characterization and biological activity. J Antibiot (Tokyo) 48:997–1003.

34. Müller S, Strack SN, Hoefler BC, Straight PD, Kearns DB, Kirby JR. 2014. Bacillaene and sporulation protect *Bacillus subtilis* from predation by *Myxococcus xanthus*. Appl Environ Microbiol 80:5603–5610.

35. May JJ, Wendrich TM, Marahiel MA. 2001. The *dhb* operon of *Bacillus subtilis* encodes the biosynthetic template for the catecholic siderophore 2,3-dihydroxybenzoate-glycine-threonine trimeric ester bacillibactin. J Biol Chem 276:7209–7217.

36. Edel-Hermann V, Lecomte C. 2019. Current status of *Fusarium oxysporum formae speciales* and races. Phytopathology 109:512–530.

37. Chen Y, Kistler HC, Ma Z. 2019. *Fusarium graminearum* trichothecene mycotoxins: Biosynthesis, regulation, and management. Annu Rev Phytopathol 57:15–39.

38. Williamson B, Tudzynski B, Tudzynski P, Van Kan JAL. 2007. *Botrytis cinerea*: The cause of grey mould disease. Mol Plant Pathol 8:561–580.

39. Abbey JA, Percival D, Abbey, Lord, Asiedu SK, Prithiviraj B, Schilder A. 2019. Biofungicides as alternative to synthetic fungicide control of grey mould (*Botrytis cinerea*) – Prospects and challenges. Biocontrol Sci Technol 29:241–262.

40. Yang H, Li X, Li X, Yu H, Shen Z. 2015. Identification of lipopeptide isoforms by MALDI-TOF-MS/MS based on the simultaneous purification of iturin, fengycin, and surfactin by RP-HPLC. Anal Bioanal Chem 407:2529–2542.

41. Ma Y, Kong Q, Qin C, Chen Y, Chen Y, Lv R, Zhou G. 2016. Identification of lipopeptides in *Bacillus megaterium* by two-step ultrafiltration and LC–ESI–MS/MS. AMB Express 6:79.

42. Kiesewalter HT, Lozano-Andrade CN, Maróti G, Snyder D, Cooper VS, Jørgensen TS, Weber T, Kovács ÁT. 2020. Complete genome sequences of 13 *Bacillus subtilis* soil isolates for studying secondary metabolite diversity. Microbiol Resour Announc 9:e01406–19.

43. Blin K, Shaw S, Steinke K, Villebro R, Ziemert N, Lee SY, Medema MH, Weber T. 2019. AntiSMASH 5.0: Updates to the secondary metabolite genome mining pipeline. Nucleic Acids Res 47:W81–W87.

44. Phelan RW, Barret M, Cotter PD, O’Connor PM, Chen R, Morrissey JP, Dobson ADW, O’Gara F, Barbosa TM. 2013. Subtilomycin: A new lantibiotic from *Bacillus subtilis* strain MMA7 isolated from the marine sponge *Haliclona simulans*. Mar Drugs 11:1878–1898.

45. Roggiani M, Dubnau D. 1993. ComA, a phosphorylated response regulator protein of *Bacillus subtilis*, binds to the promoter region of *srfA*. J Bacteriol 175:3182–3187.

46. Nakano MM, Zuber P. 1989. Cloning and characterization of *srfB*, a regulatory gene involved in surfactin production and competence in *Bacillus subtilis*. J Bacteriol 171:5347–5353.

47. Nakano MM, Xia L, Zuber P. 1991. Transcription initiation region of the *srfA* operon, which is controlled by the *comP*-*comA* signal transduction system in *Bacillus subtilis*. J Bacteriol 173:5487–5493.

48. Wang X, Luo C, Liu Y, Nie Y, Liu Y, Zhang R, Chen Z. 2010. Three non-aspartate amino acid mutations in the ComA response regulator receiver motif severely decrease surfactin production, competence development, and spore formation in *Bacillus subtilis*. J Microbiol Biotechnol 20:301–310.

49. Zeigler DR, Prágai Z, Rodriguez S, Chevreux B, Muffler A, Albert T, Bai R, Wyss M, Perkins JB. 2008. The origins of 168, W23, and other *Bacillus subtilis* legacy strains. J Bacteriol 190:6983–6995.

50. Hu F, Liu Y, Li S. 2019. Rational strain improvement for surfactin production: Enhancing the yield and generating novel structures. Microb Cell Fact 18:1–13.

51. Hamoen LW, Eshuis H, Jongbloed J, Venema G, van Sinderen D. 1995. A small gene, designated *comS*, located within the coding region of the fourth amino acid-activation domain of *srfA*, is required for competence development in *Bacillus subtilis*. Mol Microbiol 15:55–63.

52. Zhang Z, Du C, de Barsy F, Liem M, Liakopoulos A, van Wezel GP, Choi YH, Claessen D, Rozen DE. 2020. Antibiotic production in *Streptomyces* is organized by a division of labor through terminal genomic differentiation. Sci Adv 6:eaay5781.

53. Gallegos-Monterrosa R, Mhatre E, Kovács ÁT. 2016. Specific *Bacillus subtilis* 168 variants form biofilms on nutrient-rich medium. Microbiology 162:1922–1932.

54. Van Gestel J, Weissing FJ, Kuipers OP, Kovács ÁT. 2014. Density of founder cells affects spatial pattern formation and cooperation in *Bacillus subtilis* biofilms. ISME J 8:2069–2079.

55. Anagnostopoulos C, Spizizen J. 1961. Requirements for transformation in *Bacillus subtilis*. J Bacteriol 81:741–746.

56. Patrick JE, Kearns DB. 2009. Laboratory strains of *Bacillus subtilis* do not exhibit swarming motility. J Bacteriol 191:7129–7133.

57. Chen R, Guttenplan SB, Blair KM, Kearns DB. 2009. Role of the σD-dependent autolysins in *Bacillus subtilis* population heterogeneity. J Bacteriol 191:5775–5784.

58. Benoit I, van den Esker MH, Patyshakuliyeva A, Mattern DJ, Blei F, Zhou M, Dijksterhuis J, Brakhage AA, Kuipers OP, de Vries RP, Kovács ÁT. 2015. *Bacillus subtilis* attachment to *Aspergillus niger* hyphae results in mutually altered metabolism. Environ Microbiol 17:2099–2113.

59. Seemann T. 2014. Prokka: Rapid prokaryotic genome annotation. Bioinformatics 30:2068–2069.

60. Page AJ, Cummins CA, Hunt M, Wong VK, Reuter S, Holden MTG, Fookes M, Falush D, Keane JA, Parkhill J. 2015. Roary: Rapid large-scale prokaryote pan genome analysis. Bioinformatics 31:3691–3693.

61. Shen W, Le S, Li Y, Hu F. 2016. SeqKit: A cross-platform and ultrafast toolkit for FASTA/Q file manipulation. PLoS One 11:e0163962.

62. Edgar RC. 2004. MUSCLE: Multiple sequence alignment with high accuracy and high throughput. Nucleic Acids Res 32:1792–1797.

63. Waterhouse AM, Procter JB, Martin DMA, Clamp M, Barton GJ. 2009. Jalview Version 2 - A multiple sequence alignment editor and analysis workbench. Bioinformatics 25:1189–1191.

64. Wickham H. 2011. ggplot2. Wiley Interdiscip Rev Comput Stat 3:180–185.

65. Wilkins D. 2019. gggenes: Draw gene arrow maps in “ggplot2”. R package version 0.4.0.

66. Wagih O. 2017. ggseqlogo: A “ggplot2” extension for drawing publication-ready sequence logos.

67. Yu G, Zhou L, Huang H. 2020. ggmsa: Plot multiple sequence alignment using “ggplot2.”

68. Price MN, Dehal PS, Arkin AP. 2010. FastTree 2 - Approximately maximum-likelihood trees for large alignments. PLoS One 5:e9490.

69. Yu G. 2020. Using ggtree to visualize data on tree-like structures. Curr Protoc Bioinforma 69:e96.

