## Supplementary Figures for "Genomic and chemical diversity of *Bacillus subtilis* secondary metabolites against plant pathogenic fungi"

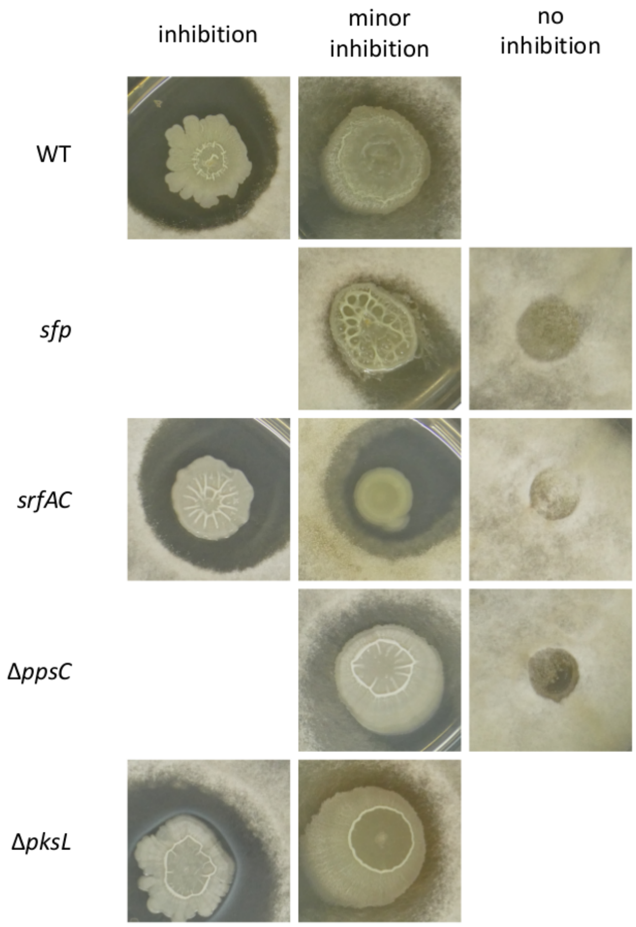


**FIG S1** The three evaluation inhibition classes, minor inhibition and no inhibition in the screening against *B. cinerea* for each *B. subtilis* genotype. No picture symbolizing the absence of the specific observation in the genotype. Minor inhibition was assigned to screening results if the fungus was inhibited but thinner hyphae were growing towards the bacterial colony.

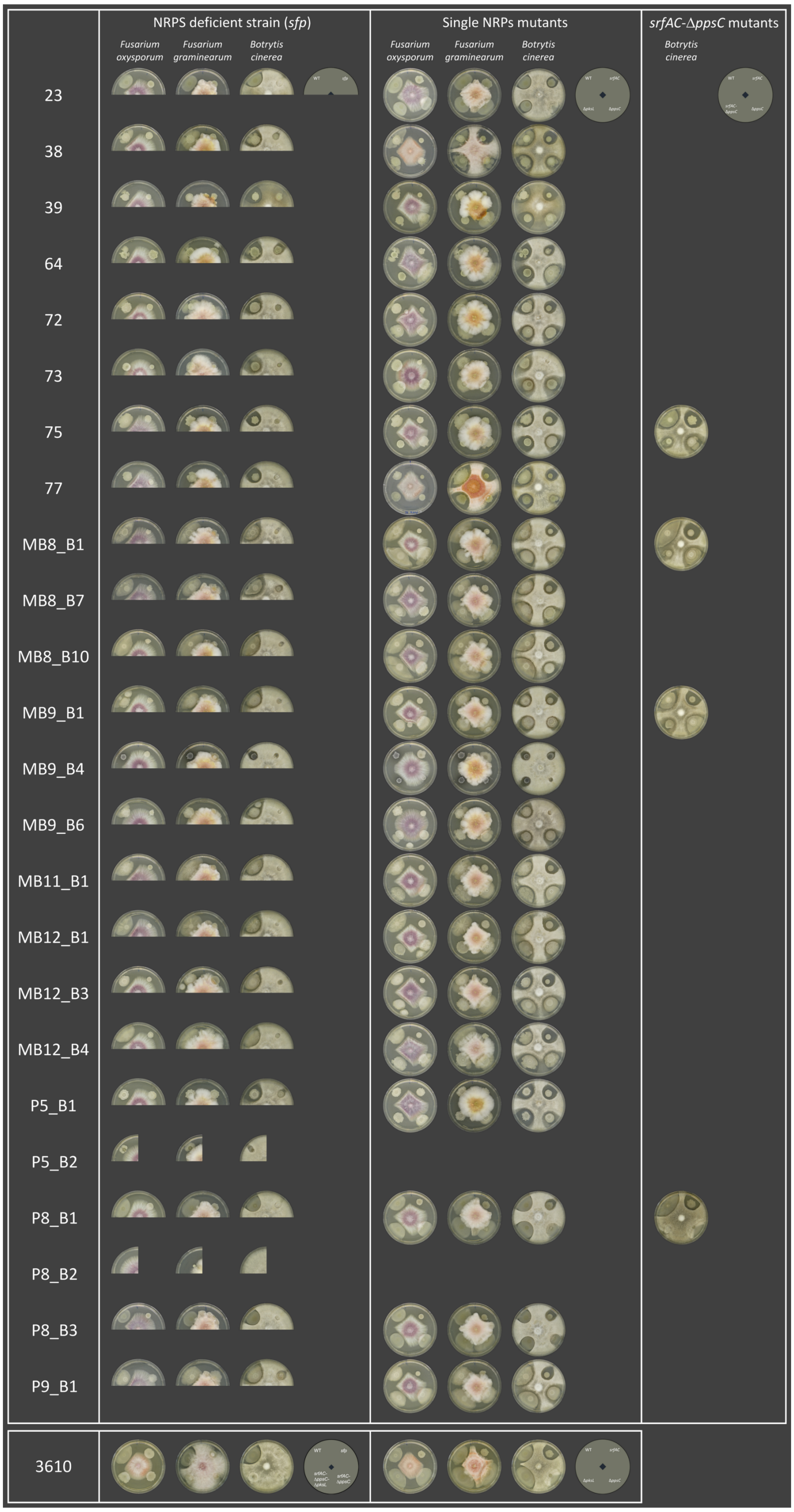


**FIG S2** Antagonism assays between the plant pathogenic fungi *F. oxysporum*, *F. graminearum* and *B. cinerea,* and the *B. subtilis* soil isolates as well as their NRP deficient *sfp* mutants (left column), single NRP mutants (middle column) and surfactin-plipastatin double mutants (right columns). The antagonism assays with *B. subtilis* 3610 are shown below. Strains were spotted as shown in the schemes on PDA plates and incubated at 21-23°C for 6 days.

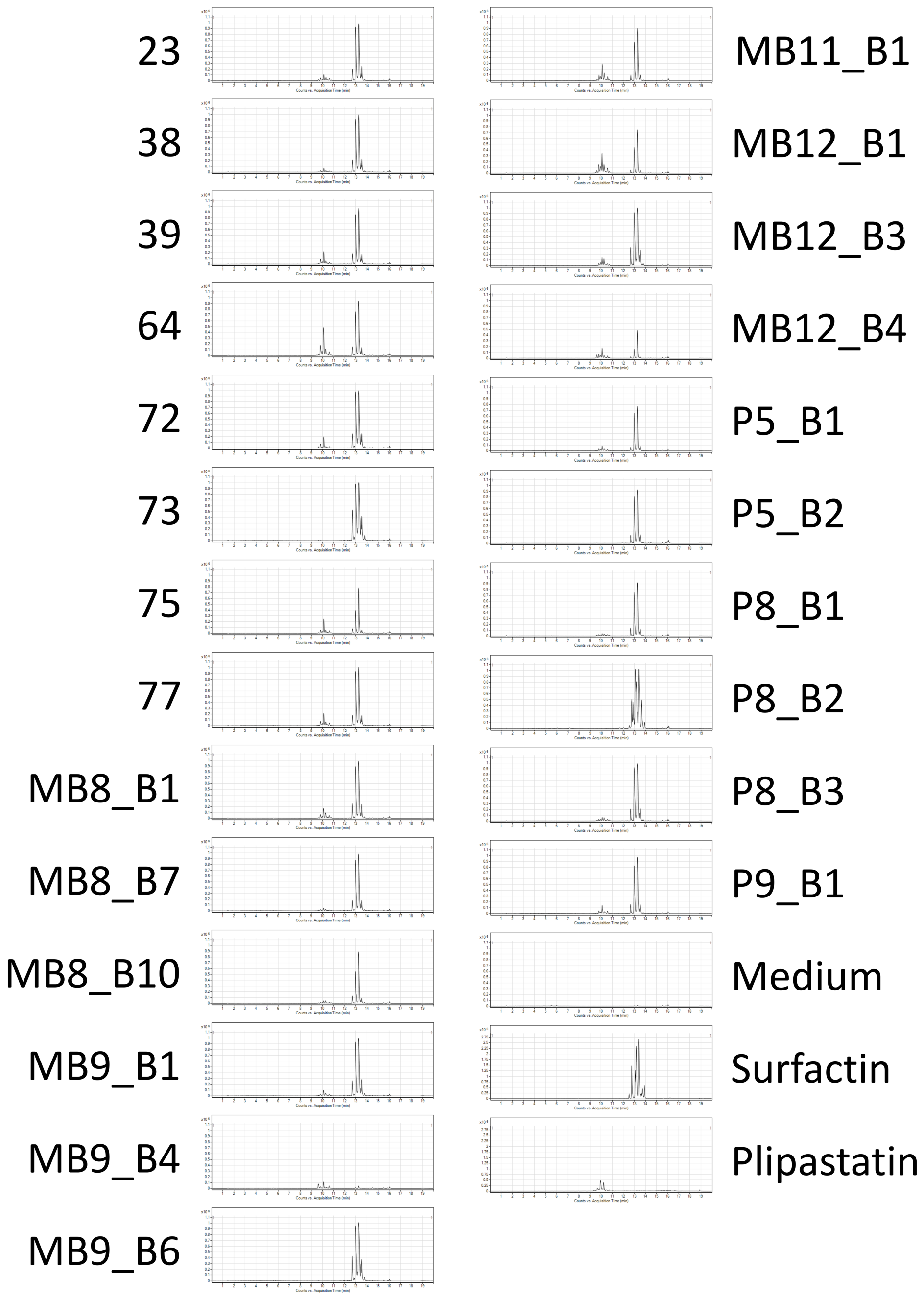


**Fig S3** Extracted ion chromatograms (m/z 1000-1600) showing the presence of surfactin and plipastatin produced by recently isolated *B. subtilis* strains grown on PDB agar medium for three days. Surfactin was detected in strain MB9_B4 but its quantity is lower compared to other isolates. Plipastatin was not detectable in extracts of strains 73, MB9_B6 and P5_B2. The chromatograms of the PDA medium and the standards of surfactin and plipastatin are shown as the last 3 panels in the second column, respectively. Surfactins, iturins and fengycins are all in the m/z range 1000–1600 that can be detected by ESI–MS.

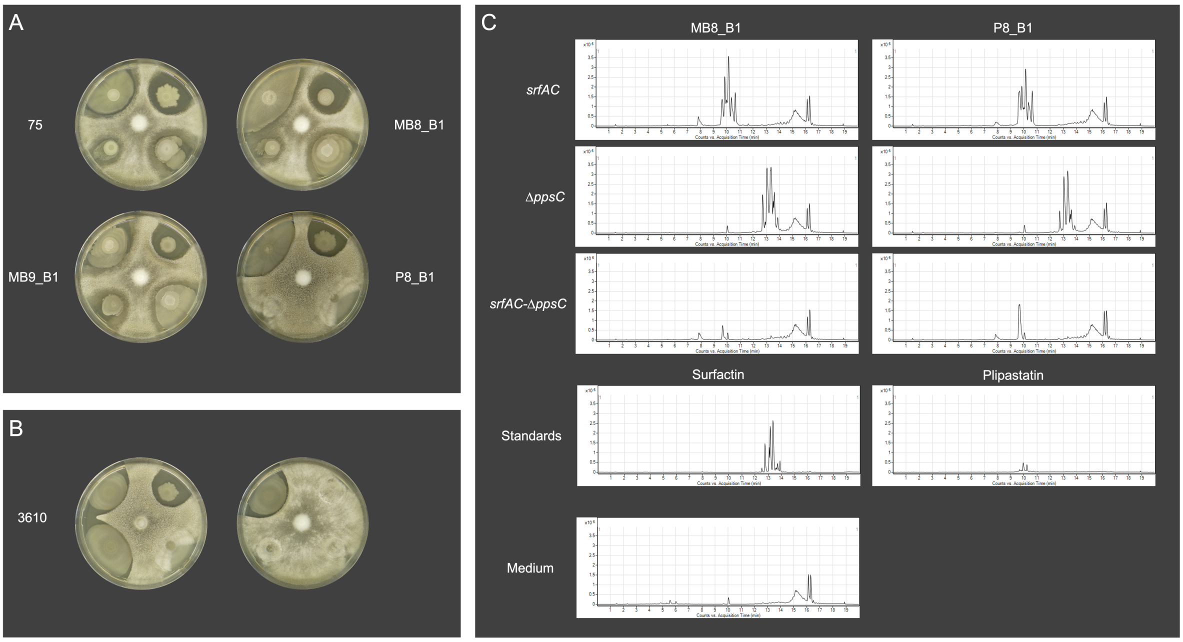


**FIG S4** (A) Antagonism assays between *Botrytis cinerea* and *B. subtilis* wild types (upper left) as well as their *srfAC* (upper right)*,* Δ*ppsC* (lower right) and *srfAC*-Δ*ppsC* (lower left) mutants. (B) Antagonism assays between *Botrytis cinerea* and *B. subtilis* 3610 as well as its NRP mutants. Left plate: WT (upper left), *srfAC* (upper right)*,* Δ*ppsC* (lower right) and Δ*pksL*(lower left). Right plate: WT (upper left), *sfp* (upper right)*, srfAC*-Δ*ppsC* (lower right) and *srfAC*-Δ*ppsC*-Δ*pksL* (lower left). Strains were co-cultivated on PDA plates and incubated at 21-23°C for 6 days. **C.** Extracted ion chromatograms (m/z 1000–1600) of *B. subtilis* mutants grown on PDA for 3 days. The chromatograms of the standards of surfactin and plipastatin and the PDA medium are shown below.

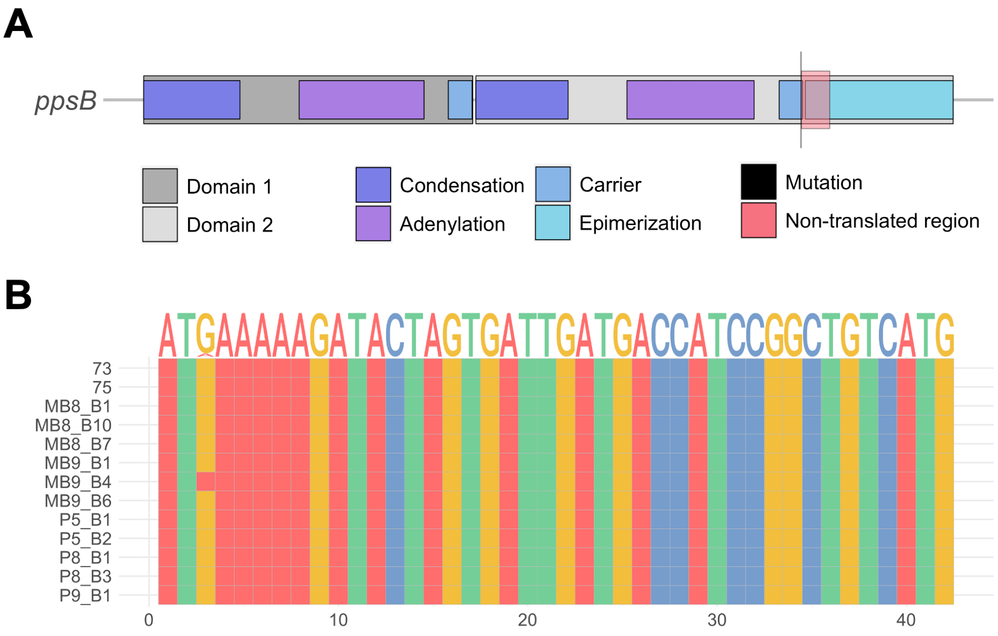


**FIG S5** (A) Strain MB9_B6 is harboring a point mutation (6224G>A), causing a point-nonsense mutation in the second domain of the *ppsB* gene due to an amino acid change from tryptophan to a termination codon (W2075X). (B) Point mutation in the *comA* gene of strain MB9_B4 (3G>A) causing a change of the translation initiating methionine.
