## Supplementary Tables for "Genomic and chemical diversity of *Bacillus subtilis* secondary metabolites against plant pathogenic fungi"

**Table S1. Bacterial strains used in this study**

| Strains | Characteristics | Reference |
|---|---|---|
| NCIB3610 | WT/Undomesticated strain | Lab stock |
| DS3337 | 3610 *sfp*::mls | (1) |
| DS1122 | 3610 *srfAC*::Tn*10* (Spec$^R$) | (2) |
| DS4085 | 3610 Δ*pksL* (Chl$^R$) | (3) |
| DS4114 | 3610 Δ*ppsC* (Tet$^R$) | (3) |
| MB8_B1 | *B. subtilis* soil isolate from sampling site 55.843861, 12.424770 | (4) |
| MB8_B7 | *B. subtilis* soil isolate from sampling site 55.843861, 12.424770 | This study |
| MB8_B10 | *B. subtilis* soil isolate from sampling site 55.843861, 12.424770 | This study |
| MB9_B1 | *B. subtilis* soil isolate from sampling site 55.843861, 12.424770 | (4) |
| MB9_B4 | *B. subtilis* soil isolate from sampling site 55.843861, 12.424770 | This study |
| MB9_B6 | *B. subtilis* soil isolate from sampling site 55.843861, 12.424770 | This study |
| MB11_B1 | *B. subtilis* soil isolate from sampling site 55.843861, 12.424770 | This study |
| MB12_B1 | *B. subtilis* soil isolate from sampling site 55.843861, 12.424770 | This study |
| MB12_B3 | *B. subtilis* soil isolate from sampling site 55.843861, 12.424770 | This study |
| MB12_B4 | *B. subtilis* soil isolate from sampling site 55.843861, 12.424770 | This study |
| P5_B1 | *B. subtilis* soil isolate from sampling site 55.788800, 12.558300 | (4) |
| P5_B2 | *B. subtilis* soil isolate from sampling site 55.788800, 12.558300 | This study |
| P8_B1 | *B. subtilis* soil isolate from sampling site 55.795200, 12.580600 | (4) |
| P8_B2 | *B. licheniformis* soil isolate from sampling site 55.795200, 12.580600 | This study |
| P8_B3 | *B. subtilis* soil isolate from sampling site 55.795200, 12.580600 | This study |
| P9_B1 | *B. subtilis* soil isolate from sampling site 55.791200, 12.575100 | (4) |
| 23 | *B. subtilis* soil isolate from sampling site 50.718551, 10.951691 | This study |
| 38 | *B. subtilis* soil isolate from sampling site 50.731996, 10.914328 | This study |
| 39 | *B. subtilis* soil isolate from sampling site 50.731996, 10.914328 | This study |
| 64 | *B. subtilis* soil isolate from sampling site 50.729170, 10.924770 | This study |
| 72 | *B. subtilis* soil isolate from sampling site 50.725876, 10.916218 | This study |
| 73 | *B. subtilis* soil isolate from sampling site 50.725876, 10.916218 | This study |
| 75 | *B. subtilis* soil isolate from sampling site 50.725876, 10.916218 | (4) |
| 77 | *B. subtilis* soil isolate from sampling site 50.725876, 10.916218 | This study |
| DTUB44 | MB8_B1 *sfp*::mls | This study |
| DTUB45 | MB8_B7 *sfp*::mls | This study |
| DTUB46 | MB8_B10 *sfp*::mls | This study |
| DTUB47 | MB9_B1 *sfp*::mls | This study |
| DTUB48 | MB9_B4 *sfp*::mls | This study |
| DTUB49 | MB9_B6 *sfp*::mls | This study |
| DTUB50 | MB11_B1 *sfp*::mls | This study |
| DTUB51 | MB12_B1 *sfp*::mls | This study |
| DTUB52 | MB12_B3 *sfp*::mls | This study |
| DTUB53 | MB12_B4 *sfp*::mls | This study |
| DTUB55 | P5_B1 *sfp*::mls | This study |
| DTUB56 | P5_B2 *sfp*::mls | This study |
| DTUB57 | P8_B1 *sfp*::mls | This study |
| DTUB58 | P8_B3 *sfp*::mls | This study |
| DTUB59 | P9_B1 *sfp*::mls | This study |
| DTUB60 | 23 *amyE*::$P_{hyperspank}$-*gfp* (Chl$^R$); *sfp*::mls | This study |
| DTUB61 | 38 *amyE*::$P_{hyperspank}$-*gfp* (Chl$^R$); *sfp*:: mls | This study |

| | | |
|---|---|---|
| DTUB62 | 39 *amyE*::P$_{hyperspank}$-*gfp* (Chl$^R$); *sfp*:: mls | This study |
| DTUB63 | 64 *amyE*::P$_{hyperspank}$-*gfp* (Chl$^R$); *sfp*:: mls | This study |
| DTUB64 | 72 *amyE*::P$_{hyperspank}$-*gfp* (Chl$^R$); *sfp*:: mls | This study |
| DTUB65 | 73 *amyE*::P$_{hyperspank}$-*gfp* (Chl$^R$); *sfp*:: mls | This study |
| DTUB66 | 75 *amyE*::P$_{hyperspank}$-*gfp* (Chl$^R$); *sfp*:: mls | This study |
| DTUB67 | 77 *amyE*::P$_{hyperspank}$-*gfp* (Chl$^R$); *sfp*:: mls | This study |
| DTUB68 | MB8_B1 *srfAC*::Tn*10* (Spec$^R$) | (4) |
| DTUB69 | MB8_B7 *srfAC*::Tn*10* (Spec$^R$) | This study |
| DTUB70 | MB8_B10 *srfAC*::Tn*10* (Spec$^R$) | This study |
| DTUB71 | MB9_B1 *srfAC*::Tn*10* (Spec$^R$) | (4) |
| DTUB72 | MB9_B4 *srfAC*::Tn*10* (Spec$^R$) | This study |
| DTUB73 | MB9_B6 *srfAC*::Tn*10* (Spec$^R$) | This study |
| DTUB74 | MB11_B1 *srfAC*::Tn*10* (Spec$^R$) | This study |
| DTUB75 | MB12_B1 *srfAC*::Tn*10* (Spec$^R$) | This study |
| DTUB76 | MB12_B3 *srfAC*::Tn*10* (Spec$^R$) | This study |
| DTUB77 | MB12_B4 *srfAC*::Tn*10* (Spec$^R$) | This study |
| DTUB79 | P5_B1 *srfAC*::Tn*10* (Spec$^R$) | (4) |
| DTUB80 | P8_B1 *srfAC*::Tn*10* (Spec$^R$) | (4) |
| DTUB81 | P8_B3 *srfAC*::Tn*10* (Spec$^R$) | This study |
| DTUB82 | P9_B1 *srfAC*::Tn*10* (Spec$^R$) | (4) |
| DTUB83 | 23 *amyE*::P$_{hyperspank}$-*gfp* (Chl$^R$); *srfAC*::Tn*10* (Spec$^R$) | This study |
| DTUB84 | 38 *amyE*::P$_{hyperspank}$-*gfp* (Chl$^R$); *srfAC*::Tn*10* (Spec$^R$) | This study |
| DTUB85 | 39 *amyE*::P$_{hyperspank}$-*gfp* (Chl$^R$); *srfAC*::Tn*10* (Spec$^R$) | This study |
| DTUB86 | 64 *amyE*::P$_{hyperspank}$-*gfp* (Chl$^R$); *srfAC*::Tn*10* (Spec$^R$) | This study |
| DTUB87 | 72 *amyE*::P$_{hyperspank}$-*gfp* (Chl$^R$); *srfAC*::Tn*10* (Spec$^R$) | This study |
| DTUB88 | 73 *amyE*::P$_{hyperspank}$-*gfp* (Chl$^R$); *srfAC*::Tn*10* (Spec$^R$) | This study |
| DTUB89 | 75 *amyE*::P$_{hyperspank}$-*gfp* (Chl$^R$); *srfAC*::Tn*10* (Spec$^R$) | (4) |
| DTUB90 | 77 *amyE*::P$_{hyperspank}$-*gfp* (Chl$^R$); *srfAC*::Tn*10* (Spec$^R$) | This study |
| DTUB91 | MB8_B1 Δ*pksL* (Chl$^R$) | This study |
| DTUB92 | MB8_B7 Δ*pksL* (Chl$^R$) | This study |
| DTUB93 | MB8_B10 Δ*pksL* (Chl$^R$) | This study |
| DTUB94 | MB9_B1 Δ*pksL* (Chl$^R$) | This study |
| DTUB95 | MB9_B4 Δ*pksL* (Chl$^R$) | This study |
| DTUB96 | MB9_B6 Δ*pksL* (Chl$^R$) | This study |
| DTUB97 | MB11_B1 Δ*pksL* (Chl$^R$) | This study |
| DTUB98 | MB12_B1 Δ*pksL* (Chl$^R$) | This study |
| DTUB99 | MB12_B3 Δ*pksL* (Chl$^R$) | This study |
| DTUB100 | MB12_B4 Δ*pksL* (Chl$^R$) | This study |
| DTUB102 | P5_B1 Δ*pksL* (Chl$^R$) | This study |
| DTUB103 | P8_B1 Δ*pksL* (Chl$^R$) | This study |
| DTUB104 | P8_B3 Δ*pksL* (Chl$^R$) | This study |
| DTUB105 | P9_B1 Δ*pksL* (Chl$^R$) | This study |
| DTUB106 | 23 *amyE*::P$_{hyperspank}$-*gfp* (Chl$^R$); Δ*pksL* (Ery$^R$) | This study |
| DTUB107 | 38 *amyE*::P$_{hyperspank}$-*gfp* (Chl$^R$); Δ*pksL* (Ery$^R$) | This study |
| DTUB108 | 39 *amyE*::P$_{hyperspank}$-*gfp* (Chl$^R$); Δ*pksL* (Ery$^R$) | This study |
| DTUB109 | 64 *amyE*::P$_{hyperspank}$-*gfp* (Chl$^R$); Δ*pksL* (Ery$^R$) | This study |
| DTUB110 | 72 *amyE*::P$_{hyperspank}$-*gfp* (Chl$^R$); Δ*pksL* (Ery$^R$) | This study |
| DTUB111 | 73 *amyE*::P$_{hyperspank}$-*gfp* (Chl$^R$); Δ*pksL* (Ery$^R$) | This study |
| DTUB112 | 75 *amyE*::P$_{hyperspank}$-*gfp* (Chl$^R$); Δ*pksL* (Ery$^R$) | This study |

| | | |
|---|---|---|
| DTUB113 | 77 *amyE*::$P_{hyperspank}$-*gfp* ($Chl^R$); Δ*pksL* ($Ery^R$) | This study |
| DTUB114 | MB8_B1 Δ*ppsC* ($Tet^R$) | This study |
| DTUB115 | MB8_B7 Δ*ppsC* ($Tet^R$) | This study |
| DTUB116 | MB8_B10 Δ*ppsC* ($Tet^R$) | This study |
| DTUB117 | MB9_B1 Δ*ppsC* ($Tet^R$) | This study |
| DTUB118 | MB9_B4 Δ*ppsC* ($Tet^R$) | This study |
| DTUB119 | MB9_B6 Δ*ppsC* ($Tet^R$) | This study |
| DTUB120 | MB11_B1 Δ*ppsC* ($Tet^R$) | This study |
| DTUB121 | MB12_B1 Δ*ppsC* ($Tet^R$) | This study |
| DTUB122 | MB12_B3 Δ*ppsC* ($Tet^R$) | This study |
| DTUB123 | MB12_B4 Δ*ppsC* ($Tet^R$) | This study |
| DTUB125 | P5_B1 Δ*ppsC* ($Tet^R$) | This study |
| DTUB126 | P8_B1 Δ*ppsC* ($Tet^R$) | This study |
| DTUB127 | P8_B3 Δ*ppsC* ($Tet^R$) | This study |
| DTUB128 | P9_B1 Δ*ppsC* ($Tet^R$) | This study |
| DTUB129 | 23 *amyE*::$P_{hyperspank}$-*gfp* ($Chl^R$); Δ*ppsC* ($Tet^R$) | This study |
| DTUB130 | 38 *amyE*::$P_{hyperspank}$-*gfp* ($Chl^R$); Δ*ppsC* ($Tet^R$) | This study |
| DTUB131 | 39 *amyE*::$P_{hyperspank}$-*gfp* ($Chl^R$); Δ*ppsC* ($Tet^R$) | This study |
| DTUB132 | 64 *amyE*::$P_{hyperspank}$-*gfp* ($Chl^R$); Δ*ppsC* ($Tet^R$) | This study |
| DTUB133 | 72 *amyE*::$P_{hyperspank}$-*gfp* ($Chl^R$); Δ*ppsC* ($Tet^R$) | This study |
| DTUB134 | 73 *amyE*::$P_{hyperspank}$-*gfp* ($Chl^R$); Δ*ppsC* ($Tet^R$) | This study |
| DTUB135 | 75 *amyE*::$P_{hyperspank}$-*gfp* ($Chl^R$); Δ*ppsC* ($Tet^R$) | This study |
| DTUB136 | 77 *amyE*::$P_{hyperspank}$-*gfp* ($Chl^R$); Δ*ppsC* ($Tet^R$) | This study |
| DTUB142 | MB8_B1 *srfAC*::Tn*10* ($Spec^R$); Δ*ppsC* ($Tet^R$) | This study |
| DTUB143 | MB9_B1 *srfAC*::Tn*10* ($Spec^R$); Δ*ppsC* ($Tet^R$) | This study |
| DTUB144 | P8_B1 *srfAC*::Tn*10* ($Spec^R$); Δ*ppsC* ($Tet^R$) | This study |
| DTUB145 | 75 *amyE*::$P_{hyperspank}$-*gfp* ($Chl^R$); *srfAC*::Tn*10* ($Spec^R$); Δ*ppsC* ($Tet^R$) | This study |

**REFERENCES**


1. Patrick JE, Kearns DB. 2009. Laboratory strains of Bacillus subtilis do not exhibit swarming motility. J Bacteriol 191:7129–7133.
2. Chen R, Guttenplan SB, Blair KM, Kearns DB. 2009. Role of the σD-dependent autolysins in Bacillus subtilis population heterogeneity. J Bacteriol 191:5775–5784.
3. Müller S, Strack SN, Hoefler BC, Straight PD, Kearns DB, Kirby JR. 2014. Bacillaene and sporulation protect bacillus subtilis from predation by myxococcus xanthus. Appl Environ Microbiol 80:5603–5610.
4. Thérien M, Kiesewalter HT, Auria E, Charron-Lamoureux V, Wibowo M, Maróti G, Kovács ÁT, Beauregard PB. 2020. Surfactin production is not essential for pellicle and root-associated biofilm development of Bacillus subtilis. Biofilm 2:100021.

**Table S2. Fungal strains used in this study**

| Fungal species | Isolation | Source |
|---|---|---|
| *Fusarium oxysporum* IBT 40872 | From surface of an equipment in a Danish factory, 2005, by Anne Svendsen | IBT Culture Collection of Fungi, DTU |
| *Fusarium graminearum* IBT 41925 | From malting barley, Schackenborg, Denmark, 2012, by Ulf Thrane | IBT Culture Collection of Fungi, DTU |
| *Botrytis cinerea* IBT 42565 | From indoor air in a Kitchen, Kongens Lyngby, Denmark, 2018, by Birgitte Andersen | IBT Culture Collection of Fungi, DTU |
